# Lateral mechanical impedance rather than frontal, promotes cortical expansion of roots

**DOI:** 10.1101/860155

**Authors:** Xuanjun Feng, Jing Xiong, Yue Hu, Liteng Pan, Zhengqiao Liao, Xuemei Zhang, Wei Guo, Fengkai Wu, Jie Xu, Erliang Hu, Hai Lan, Yanli Lu

**Affiliations:** State Key Laboratory of Crop Gene Exploration and Utilization in Southwest China, Sichuan Agricultural University. Wenjiang, Sichuan, China; Maize Research Institute, Sichuan Agricultural University, Wenjiang, Sichuan, China; Key Laboratory of Biology and Genetic Improvement of Maize in Southwest Region, Ministry of Agriculture, China

## Abstract

It has long been considered that mechanical impedance on root will restrict root elongation and consequently promote radial growth. However, we did observe radial expansion but not elongation restriction in maize seedlings after short growth in sands. Mechanical impedance of soil can be classified into frontal- and lateral-type based on the interaction site of root. Therefore, we suspected that radial expansion might be mainly stimulated by lateral rather than frontal impedance. To verify our speculation, frontal and lateral impedance was provided separately. Small plastic caps were used to provide unique frontal impedance on root tip and cylindrical plastic containers were used to provide lateral impedance. Plastic caps could reduce root length remarkably. However, the radial expansion of plastic-cap-fitted roots was significantly inferior to that of the sand-cultured roots. Microstructural analysis revealed that sand-condition thickened root largely dependents on cortical expansion, whereas plastic cap did it mainly by thickening stele. In cylindrical plastic containers, mechanical impedance came only from the lateral direction and promoted the expansion of cortex just as sand-condition. Thus, we proposed that the expansion of the cortex and the consequent radial growth is mainly due to lateral impedance when growing in sands.

## Introduction

The growth of roots in soil can be limited by the physical, chemical, and biological properties of soil, among which physical properties have been reported to be most strongly linked to root elongation [1–3]. In terms of the physical limitations on root growth, mechanical impedance (caused by soil that is too compact to enable rapid root penetrance) mainly affects root systems by hindering root elongation and concurrently promoting root thickening [1,3,4]. Soil is a dense medium, and it is necessary for root tips to generate sufficiently mechanical force to penetrate through the soil in the absence of continuous pores of a sufficiently large diameter [1,5]. Intuitively, thin roots, given their cross section, may penetrate soil easier as they encounter less resistance than do thicker roots. Moreover, thin roots will reorient their growth easily to circumnavigate the obstacle [1,5]. Therefore, it sounds contradictory that roots grow thicker in response to mechanical impedance [1,4,6]. Indeed, previous experimental evidences related to the effect of root diameter on penetration resistance were often contradictory [1,7]. But recently, more studies on different plant species have indicated that thicker roots rather than thinner roots can more effectively penetrate compact soil layers [4,8].

Actually, mechanical impedance of soil can be classified into frontal- and lateral-type based on the position of root, and frictional (lateral) impedance between roots and the soil may account for up to 80% of total mechanical impedance [4,9]. All previous studies on mechanical impedance have obtained data from plants in which the entire root system has been embedded in various substrates, such as glass beads, soil, sands, and phytagel [1,3–5,10]. In these studies, it was not possible to distinguish the effects of frontal and lateral impedance. So, It is not entirely clear as to whether frontal mechanical impedance acts as a direct cue to restrict root elongation and promote radial expansion. In this study, the objective was to determine which of the frontal- or lateral- impedance was the main stimuli on radial expansion of root.

## Materials and methods

### Plant materials and growth conditions

To produce the seedlings used in this study, we germinated maize seeds in rolled-up germinating test paper, so as to generate seedlings with relatively straight roots of approximately 3 to 5 cm in length. These seedlings were then subjected to different growth conditions (water-, sand-, semi-sand and cap-fitted-conditions). Silica sand with a particle size about 2 mm in diameter was used for sand and semi-sand conditions. The maize seedlings were grown in a greenhouse at approximately 26°C under a 14-h light/10-h dark photoperiod. Schematic diagrams of plastic cap and cylindrical plastic container were showed in Figure 1a and 1b, respectively. Plastic cap was used for cap-fitted condition. It was prepared in the following manner: a rubber tube with a 3 mm inner diameter was cut into 1 cm lengths and one end of the 1 cm rubber tube was blocked. Then, a tiny elastic thread was fixed to two flanks of the plastic cap. When used, plastic cap was fitted on root tip and elastic thread was attached to the raft where seedlings were anchored (Fig 1D). The cylindrical plastic container was used for semi-sand condition. It was prepared in the following manner: first, plastic pipe with a 20 mm inner diameter was cut into 5 cm lengths; second, one end of the 5 cm plastic pipe was blocked by a plastic sheet and scotch tape; at last, making a hole in the center of the plastic sheet about 3 mm. When used, root of seedling was buried in the cylindrical plastic container with sands and root tip was exposed to the outside through a hole (Fig 1E). Four kinds of growth conditions were showed in Fig 1.

**Fig 1.**
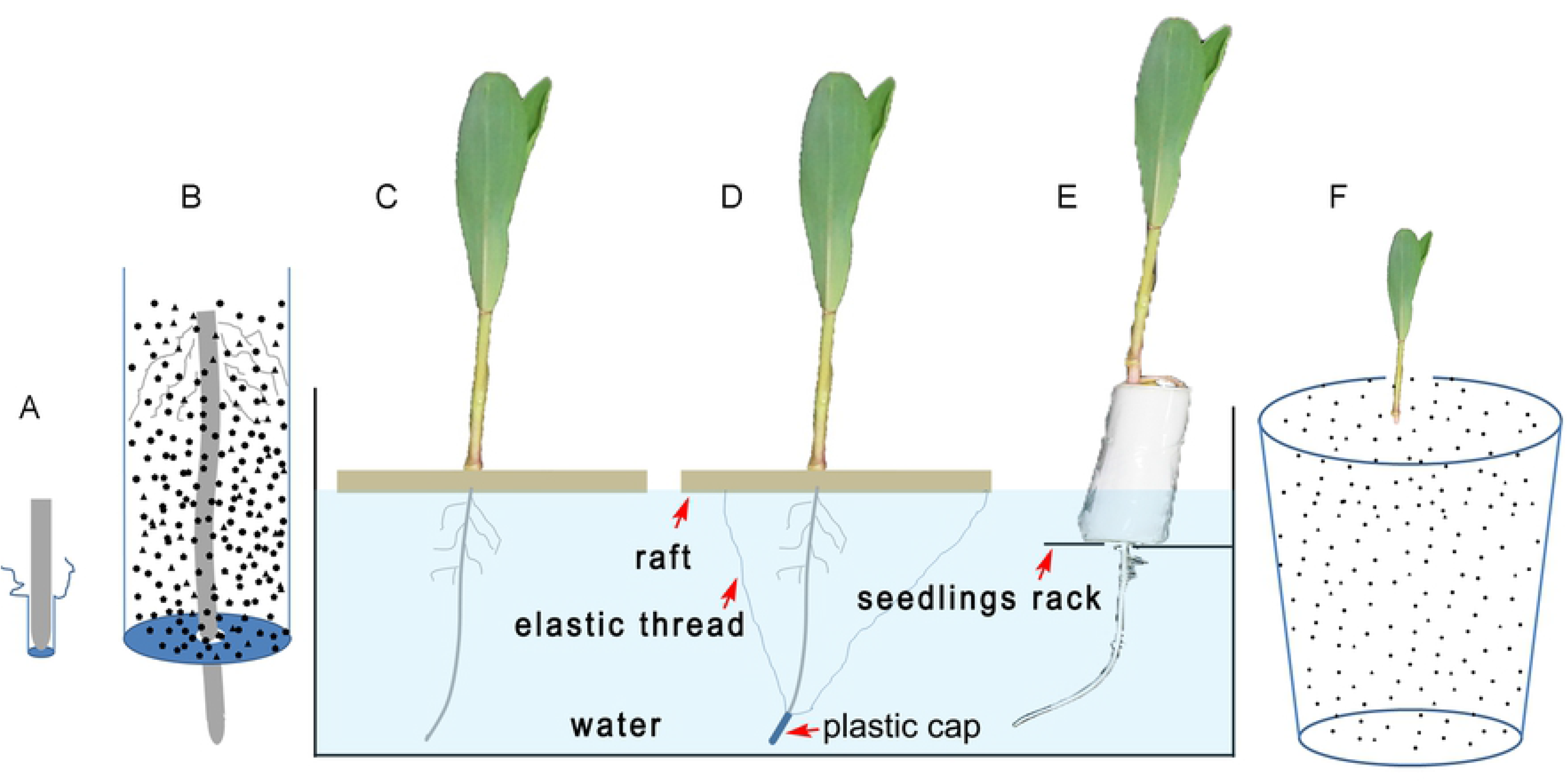
Schematic diagram of treatment conditions. (A) Root tip in plastic cap. (B) root in cylindrical plastic container. It is water-, cap-fitted-, semi-sand and sand-condition from (C) to (F) successively. Black points in (B) and (F) represent silica sands.

### Preparation of paraffin sections

Segments of root from the root tip and the region (1 cm in length) near the root–hypocotyl junction were collected from three biological replications (5 seedlings were used in each replication) and stored in Carnoy’s fixative (ethanol: glacial acetic acid = 3:1) at room temperature for 12 h. The samples were then subjected to an ethanol gradient, consisting of 75% ethanol for 4 h, 85% ethanol for 4 h, 100% ethanol overnight, and 100% ethanol + basic fuchsin for 4 h. Thereafter, the samples were transferred to xylene (two 1 h treatments). The subsequently obtained transparent samples were embedded in melted paraffin for 12 h, after which the paraffin was replaced with new paraffin and saturated for more than 24 h. The samples were then embedded in paraffin using plastic molds. Cross-sections of the embedded roots (with a thickness of approx. 15 µm) were prepared using an RM2255 microtome (Leica, Germany). For each replication, 10 slides (2 slides per sample) were prepared and then stained with 1% toluidine blue for 1 min, followed by three washes in double-distilled water. After inspection for breakage and uniformity, two sections of each slide were selected for image analysis.

### Image acquisition and analysis

For the investigation of cell length, fresh primary root tips (approx. 0.8 cm in length) were longitudinally sliced by hand and stained with 50 µg/mL propidium iodide for 10 min. Then, the samples were washed twice and observed using a Zeiss LSM 800 confocal laser scanning microscope (Germany). The view field was located up to 400 µm from the root tip border, and images were acquired using an objective lens with a 10x magnification. To obtain the relative full view of the roots after propidium iodide staining, 20 lateral roots of each sample were selected and their images were acquired using the Tiles model of Zeiss LSM 800. Micrographs of sections were acquired using an Olympus microscope (IX73, Japan) fitted with a DP80 digital camera under white light using a 10x objective lens. Images of root hairs were obtained using an Olympus anatomical lens (SZX2-ILLT, Japan) fitted with a DP71 digital camera under white light using a zoom magnification of 6.4x with its 1x objective. Three biological replications were established, 5 seedlings of each replication were used for cell length analysis and 10 seedlings of each replication were used for the analyses of root length and diameter. Images were analyzed for anatomical traits using Image J software. Cross-sectional diameter and stele diameter were measured directly, whereas cortex thickness was determined by subtracting the stele diameter from the cross- sectional diameter. The difference between samples was based on the Student’s t-test.

### RNA extraction and PCR

To investigate the expression of *Cyclin* genes, which are the marker genes reflecting cell division activity [11, 12], total RNA was isolated from root tips using the TRIzol reagent (Invitrogen) and treated with RNase-free DNase I (Takara). An iScript™ cDNA Synthesis Kit (Bio-Rad) was used to synthesize cDNA, using 3 µg of total RNA from each sample. Subsequently, quantitative real-time reverse transcription PCR (qRT-PCR) was performed using SsoFast™ EvaGreen® Supermix (Bio-Rad) in 96-well optical reaction plates in a Bio-Rad CFX96 Touch™ thermal cycler. Data were processed as described in our previous publication [13]. Quantification was performed using the 2−∆∆Ct method [14], where ∆∆Ct is the difference in threshold cycles between specific genes and the reference housekeeping gene eF1a. The primers used for amplification are listed in Table 1.

**Table 1.**
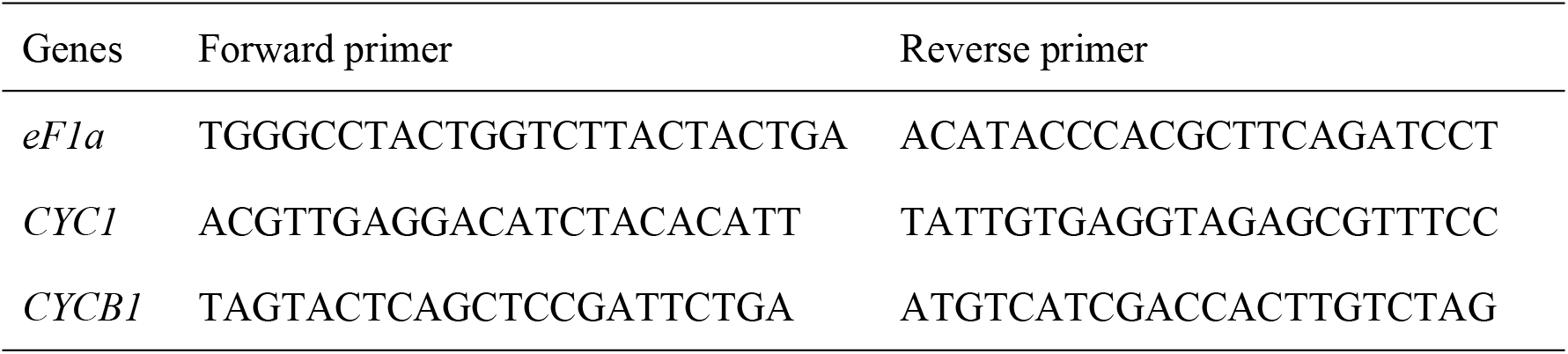
Primers used for quantitative real-time reverse transcription PCR (qRT-PCR).

## Results

### Frontal impedance reduced root length by restricting cell division and elongation

At the beginning, maize seedlings were exposed to sand-condition for 3 and 6 days to investigate the effect of sands provided mechanical impedance. Water cultured seedlings were used as controls. When compared to controls, 3 days’ sand-cultured seedlings did not display obvious changes in root length, but we did observe a substantial expansion of root diameter, particularly in the region within 3 cm of the root apex, which was newly developed after transplant (Fig 2A, 2B). However, by prolonging growth time to 6 days, we found a significant reduction of root length under sand-condition (Fig 2A). From this result, it was believable that elongation restriction and radial expansion were not always accompanying. Thereafter, 3 days of treatment was chose. Next, cap-fitted-condition was added and seedlings were treated for 3 days. Small plastic caps were fitted onto root tips to provide unique frontal impedance. After 3 days of growth, root length, cell length, expression of *Cyclin* genes (marker genes reflecting cell division activity) and root diameter, were investigated [11,12]. We found that in plants with plastic caps, root length was reduced by one-third when compared with that of controls. There was, however, no statistically significant difference in root length between the sand-condition seedlings and controls (Fig 2C). At the microscopic and molecular level, we observed that the presence of plastic cap promoted a marked reduction in cell length and down-regulated expression of *cyclin B* (Fig 2D, 2E), thereby implying low cell division activity under frontal impedance. Accordingly, we could deduce that reductions in cell division and cell length contributed to determining a shorter root length under frontal impedance. In addition, we observed that a larger number of root hairs had emerged on the root of sand-condition seedlings than in the cap-fitted seedlings and controls (Fig 2F).

**Fig 2.**
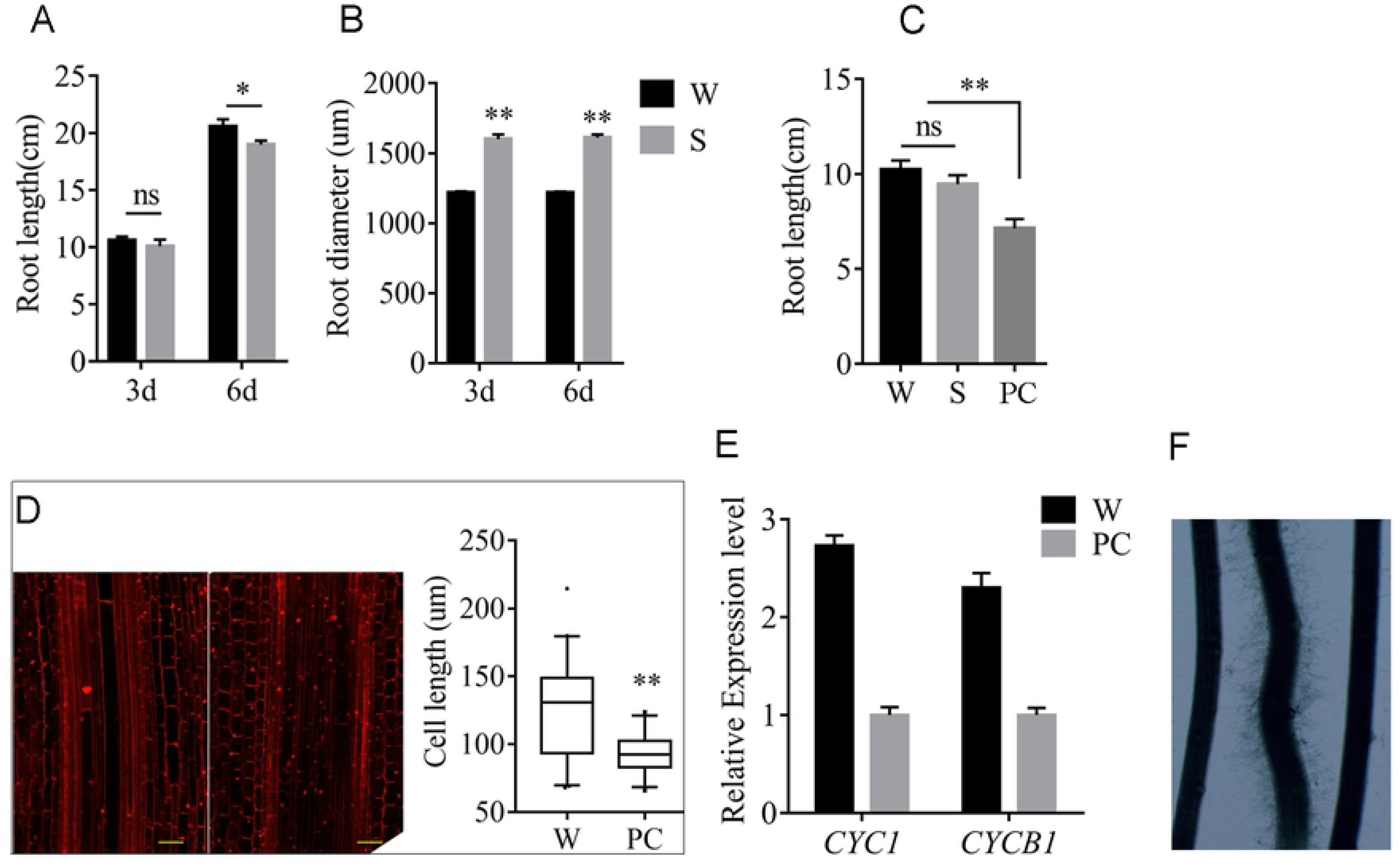
Frontal impedance reduced root length by restricting cell division and elongation. Root length (A) and diameter (B) after 3 or 6 days of growth in water-(W) and sand- (S) conditions. (C) Root length after 3 days of growth under W, S, and plastic-cap-fitted (PC) conditions. (D) Confocal micrographs of primary root tips after propidium iodide (PI) staining (left: W, right: PC) and cell length after 3 days of growth under W and PC conditions. The yellow scale bars represent 100 µm. (E) Expression level of two *Cyclin* genes (cell division marker genes) and *eF1a* as an internal control. (F) Root hairs on primary root viewed under an anatomical lens (from left to right: W, S, and PC). **P < 0.01; *P < 0.05; ns: non-significant.

### Frontal impedance increased stele diameter but not cortical expansion

As to root diameter, significant differences were observed between three kinds of seedlings (Fig 3, 2C). Notably, the length of plastic-cap-fitted roots was reduced by one-third compared with those grown in sand, although the root diameter was thinner in the cap-fited roots (Fig 3A, 3B). This finding was unexpected and implies that simple frontal impedance has a slight facilitating effect on radial expansion.Furthermore, root tips were sliced using a microtome for microexamination. The roots of sand-condition seedlings displayed remarkable cortical cell expansion compared with the roots of controls and plastic-cap-fitted roots, and there was no difference between controls and plastic-cap-fitted roots (Fig 3A, 3C). In terms of stele diameter, plastic-cap-fitted roots were thicker than both water- and sand-condition roots, whereas there was no difference between water- and sand-condition roots (Fig 3A, 3D). These observations indicate that simple frontal impedance did not induce cortical expansion, but it did increase the stele diameter of roots.

**Fig 3.**
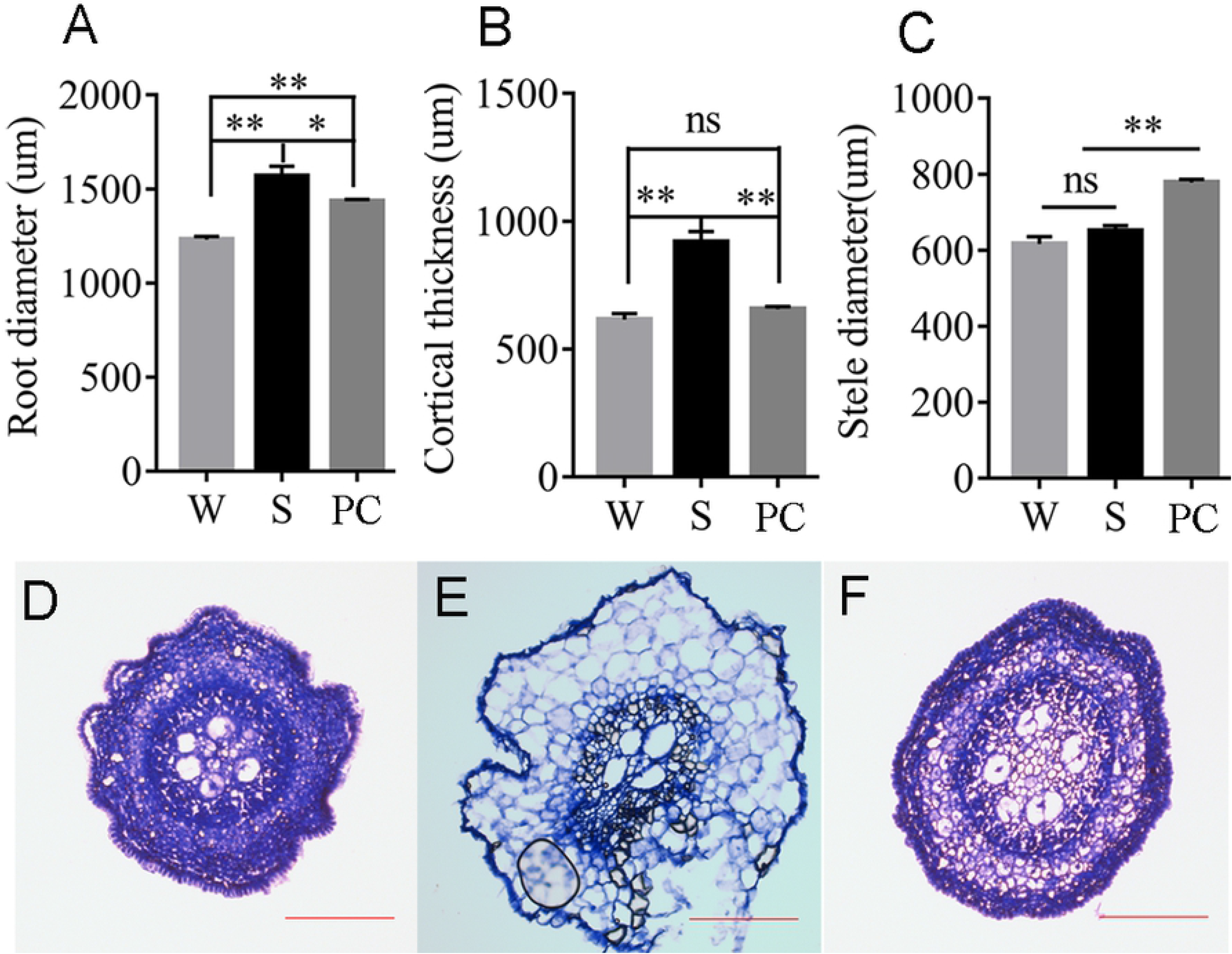
Root diameter (A), cortical thickness (B) and stele diameter (C) of root tips. Micrographs of a cross-section under water-, sand- and plastic-cap-fitted conditions from (D) to (F) successively. The red scale bars represent 100 µm. **P < 0.01; *P < 0.05. W, S and PC represent water-, sand-, and plastic-cap-fitted conditions, respectively.

### Lateral impedance promoted cortical cell expansion

Given that frontal impedance conferred by the presence of a plastic cap did not induce notable cortical expansion, the effect of lateral impedance on cortex was examined. Cylindrical plastic containers were used to provide semi-sand-cultured conditions (shown in Fig 4A), in which mechanical impedance was imposed only in a lateral direction. After 3 days of treatment, root diameter and microstructure were analyzed using the mature region (2 cm below the root–hypocotyl junction). The results showed that both sand and semi-sand culturing promoted cortical expansion when compared with the roots of water-condition (Fig 4B, 4D). However, the increment of cortical expansion at this region was smaller than that in the apical part of roots grown in sand-condition (Fig 3B, 4C). This difference could be attributable to that root tip is a newly developed tissue sensing de novo stimuli of lateral impedance after transplant.

**Fig 4.**
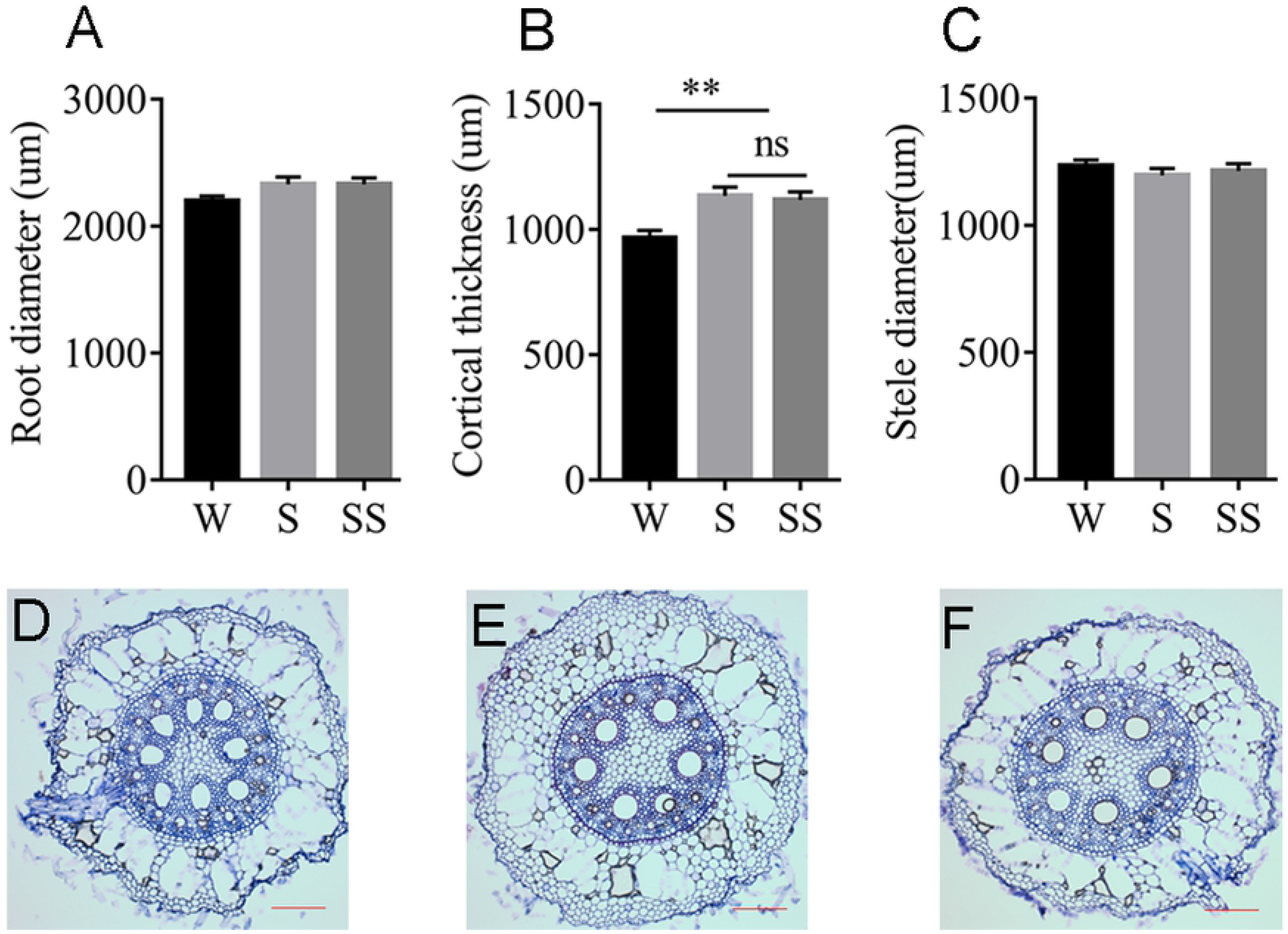
Root diameter (A), cortical thickness (B) and stele diameter (C) of mature region. Micrographs of cross-sections under water-, sand- and semi-sand- conditions from (D) to (F) successively. The red scale bars represent 100 µm. **P < 0.01. W, S and SS represent water-, sand- and semi-sand- conditions, respectively.

## Discussion

Considerable evidences have amassed to indicate that mechanical impedance can reduce the elongation and promote the radial expansion of roots [1,4,5,10]. Nevertheless, all previous studies on mechanical impedance have obtained data from plants in which the entire root system has been embedded by various substrates, such as glass beads, soil, sand, and phytagel [1,3–5,10]. In these studies, it was not possible to distinguish the effects of frontal and lateral impedance, and consequently, root shortening and thickening always appeared concurrently. In the present study, plastic caps were used to provide simple frontal impedance on the root tips. Subsequent microstructural observations and expression analysis of cell division marker genes indicated that reduced cell division and cell length jointly determined the short root lengths observed under frontal impedance. However, although the presence of plastic caps resulted in seedlings with shorter roots than those of seedlings cultured in sand, we found that the diameter of plastic-cap-fitted roots was smaller than that of sand-condition roots, implying that frontal impedance has a slight facilitating effect on radial expansion. Mechanical impedance has been demonstrated to increase root diameter mainly via the expansion of cortical cells [4,15]. Based on our results, however, simple frontal impedance did not induce the expansion of the cortex, whereas it did increase the diameter of the stele. Thus, we speculate that the cues promoting cortical expansion may arise from lateral impedance, such as frictional impedance. Actually, the frictional resistance between roots and the soil has been reported to account for up to 80% of the total mechanical impedance [4,9]. Vertical pressure is, however, considered an unlikely candidate, because it has previously been reported to constrict radial expansion and compress cortical cells [3,16].

The expansion of cortical cells is controlled by the orientation of cortical microtubules or actin microfilaments and the consequent deposition of cellulose microfibrils [16–18]. This process is also associated with the secretion of mucilage, for instance, the treatment of root cap cells with actin microfilament inhibitor was found to reduce mucilage production, while disorganization of cortical microtubules has been observed to impair the release of mucilage from seed coat secretory cells [17,19]. Thus, cortical cell expansion appears to be associated with the process of mucilage production and consequently overcomes frictional resistance between roots and sand particles to promote root penetration in sands.

## Conclusions

We propose that cortical expansion and the consequent thickening of roots are primarily induced by lateral impedance (friction) rather than frontal impedance when roots grow in sands.

## Acknowledgments

We thank Prof. Xuejun Hua for his edit and suggestion. This work was supported by the National Natural Science Foundation of China (31801371), the National Ten Thousands Program for Young Top Talent and the International Cooperative Projects of Sichuan province, China (2017HH0027).

